# Phenotype–genotype discordance in antimicrobial resistance profiles of Gram-negative uropathogens recovered from catheter-associated urinary tract infections in Egypt

**DOI:** 10.1101/2025.04.17.649370

**Authors:** Mohamed Eladawy, Nathan Heslop, David Negus, Jonathan C. Thomas, Lesley Hoyles

## Abstract

Catheter-associated urinary tract infections (CAUTIs) are among the most common healthcare-associated infections in low- and middle-income countries (LMICs). In this study, phenotypic (EUCAST AMR profiles) and genomic data were generated for 132 isolates (67 *Escherichia coli*, 14 *Pseudomonas aeruginosa*, 11 *Klebsiella pneumoniae*, 9 *Proteus mirabilis*, 8 *Providencia* spp., 5 *Enterobacter hormaechei*, and 18 rare uropathogens) recovered from CAUTIs in an Egyptian hospital. Among the *Escherichia coli* isolates, phylogroup B2 was most represented (53.7 %), followed by B1 (19.4 %), A (11.9 %), D (7.4 %), F (5.9 %) and C (1.4 %). Several (22/132, 16.6 %) isolates were multidrug-resistant, while almost half (62/132, 46.9 %) were extensively drug-resistant. Comparison of phenotypic data with genotypic data from three different AMR-profiling tools (ResFinder, CARD, AMRFinder) highlighted phenotype–genotype discordance as an important consideration in resistome studies in Egypt, with a total concordance of 91.1 %, 85.7 %, and 80.3 % for ResFinder, CARD and AMRFinder, respectively. *Pseudomonas*, at the species level, exhibited the greatest discordance. At the antimicrobial level, meropenem was subject to greatest discordance. In addition to the findings from our comparative analyses, new AMR variants are reported for Egypt for *Pseudomonas* (*OXA-486, OXA-488, OXA-905, IMP-43, PDC-35, PDC-45, PDC-201*) and *Escherichia coli* (*TEM-176, TEM-190*). In summary, we show that there is phenotype–genotype discordance in AMR profiling among CAUTI isolates, highlighting the need for comprehensive approaches in resistome studies. We also show the genomic diversity of Gram-negative uropathogens contributing to disease burden in a little-studied LMIC setting.

## INTRODUCTION

Urinary tract infections (UTIs) are an increasing public health problem caused by a range of uropathogens. The frequency of healthcare-associated UTIs is 12.9 %, 19.6 % and 24 %, respectively, in the United States, Europe and developing countries ^1^. In 2019, bacterial antimicrobial resistance (AMR) contributed to 48,700 deaths associated with UTIs out of 541,000 cases of infectious syndromes ^2^. Resistance to almost all antibiotics in healthcare-associated UTIs is above 20 %, with significant geographical variation ^1^. However, meta-analysis and surveillance data are scarce from low- and middle-income countries (LMICs) ^3, 4^. Egypt initiated a national surveillance program for healthcare-associated infections (HAIs) with international collaborators and national entities to establish benchmarks to outline the microbiological profiles and reduce AMR in clinical settings. Phase one of the National Action Plan showed nearly 50 % of *Escherichia* (*Esc*.) *coli* and 75 % of *Klebsiella pneumoniae* isolates from urine were resistant to quinolones and cephalosporins, respectively ^5^. Across 27 countries, Egypt came in third place for total consumption of antibacterials (1355.4 tonnes) in 2020 ^6^.

In Egypt, catheter-associated UTIs (CAUTIs) are among the most common types of HAIs affecting patients in intensive care units (ICUs) ^7-10^. Between 2011 and 2012, 15 % (69/472) of HAIs were UTIs and the rate of CAUTIs was 1.2 per 1,000 device-days ^7^. Another surveillance showed that UTIs represented 15 % (406/2688) of HAIs, the rate of CAUTI development increased up to 1.9/1,000 device-days and CAUTIs represented 98.3 % of UTIs ^8^. An epidemiological study (between 2011 and 2017) of carbapenem-resistant *Enterobacteriaceae* in Egyptian ICUs showed that UTIs accounted for 14.8 % (236/1598) of HAIs, with 63.8 % of UTIs catheter-related ^10^. The second most common HAIs were UTIs (12.2 %) in Egyptian clinical settings ^9^.

There are few reports on the genomic diversity of uropathogens found in Egyptian clinical settings ^11-16^. However, high rates of AMR are observed in Egypt beside documented transmission of carbapenem-resistant strains from Egypt to other countries ^17-19^. We collected a range of Gram-negative uropathogens (n = 132 isolates) from an Egyptian hospital over a 9-month period, and generated genotypic and phenotypic data for them. Our aim was to investigate the AMR patterns associated with UTIs in an Egyptian clinical setting and provide evidence that could inform the development of effective antimicrobial surveillance plans and targeted infection control policies. In addition, we assessed the agreement/concordance between phenotypic and genotypic data to evaluate the usefulness of such data from a diagnostic perspective.

## METHODS

### Recovery of isolates

Anonymized isolates (n = 132) with no patient data were recovered from urinary catheters between December 2020 and August 2021 by staff at the Urology and Nephrology Center, Mansoura, Egypt during routine diagnostic procedures as described previously ^16^. The study of anonymized clinical isolates beyond the diagnostic requirement was approved by the Urology and Nephrology Center, Mansoura, Egypt. No other ethical approval was required for the use of the clinical isolates.

### Antimicrobial susceptibility testing

Antimicrobial susceptibility testing was performed using the disc diffusion test (DDT) on Mueller-Hinton agar (Oxoid Ltd, UK) ^16^. Twelve different antimicrobial discs (Oxoid Ltd, UK) were used: penicillins – piperacillin/tazobactam (30/6 μg); cephalosporins – ceftazidime (10 μg), cefepime (30 μg); monobactams – aztreonam (30 μg); carbapenems – doripenem (10 μg), meropenem (10 μg); aminoglycosides – tobramycin (10 μg), amikacin (30 μg); quinolones – ciprofloxacin (5 μg), levofloxacin (5 μg); miscellaneous – nitrofurantoin (100 μg), sulfamethoxazole/trimethoprim (1.25–23.75 μg).

Results were interpreted according to breakpoints of EUCAST guidelines version 12.0, 2022 (http://www.eucast.org/clinical_breakpoints/), with EUCAST reference strains - *Esc. coli* ATCC 25922 and *Pse. aeruginosa* ATCC 27853 - used for quality control purposes.

### DNA extraction and whole-genome sequencing

DNA was extracted from isolates and their genomes were sequenced and assembled as described previously ^16, 20^.

### Bioinformatic analyses

Bakta v1.5.1 (database v4.0) was used for annotating genes within genomes ^21^. sourmash v4.8.4 ^22^ was used to compare and identify (with Genome Taxonomy Database release 214) species for all genomic data (**Supplementary Table 1**). Genomes that could not be assigned to a species by sourmash were identified using ribosomal multilocus sequence type (rMLST) (https://pubmlst.org/species-id; ^23^), and by comparison with relevant reference sequences (**Supplementary Table 1**) using FastANI v1.33 ^24^.

Serotypes were determined using ECTyper v1.0.0 (database v1.0) ^25^. Phylogrouping of *Esc. coli* was achieved using Ezclermont v0.7.0 ^26^. MLST of each isolate was determined using *mlst* v2.23.0 (https://github.com/tseemann/mlst) and relevant schema from http://pubmlst.org ^27-33^.

AMR genes were identified using (accessed February 2025) the Resistance Gene Identifier v6.0.3 of the Comprehensive Antibiotic Resistance Database (CARD) v4.0.0 ^34^, ResFinder v4.6.0 (ResFinder database v2.4.0, PointFinder database v4.1.1) ^35^, and AMRFinder v4.0.3 (database version 2024-12-18.1) ^36^. Genes responsible for efflux pumps were excluded from this study ^37^. Only AMR genes that showed perfect or strict matches (identity ≥ 90 %) were reported.

For the *Esc. coli* genomic data, a phylogenetic tree was constructed using core genes identified using Panaroo v1.3.4 ^38^ (strict mode, identity threshold 98 %, alignment option “core”). Core genes were present in all 67 *Esc. coli*, and were aligned with prank (v170427) ^39^. Gene alignments were concatenated with FASconCAT-G (v1.02) into a single alignment ^40^. Phylogenetic analysis was performed using the concatenated sequences and RaxML v1.2.0 (GTR+G8+IO) ^41^. Node support was determined using 1000 non-parametric bootstrap replicates. The phylogenetic tree was visualized using iTOL v6.9 ^42^. Virulence traits encoded in genomes were detected via BLASTP (≥90 % coverage, ≥70 % identity) with the core protein sequence dataset from the Virulence Factors Database (VFDB) ^43^.

### Comparison of AMR genotypes and phenotypes

According to EUCAST guidelines, “*susceptible, standard dosing regimen*” (S) and “*susceptible, increased exposure*” (I) categories are grouped under the term “*susceptible*”. Using CARD, ResFinder, and AMRFinder, genomic data of isolates were compared with DDT results for 132 isolates against 12 antimicrobials [except when certain species lacked breakpoints; i.e. nitrofurantoin was used only for *Esc. coli* (n = 67), sulfamethoxazole/trimethoprim was used only for *Enterobacterales* (n = 114)]. For each combination, concordance was considered positive if a) WGS data were predicted to encode AMR genes and the isolate had a phenotypic resistant profile “true resistant” (WGS-R/DDT-R) or b) WGS data were not predicted to encode AMR genes and the isolate had a phenotypic susceptible profile “true susceptible” (WGS-S/DDT-S) ^44, 45^. Discordance was considered positive where major errors (MEs) or very major errors (VMEs) were detected. MEs (i.e. false positives; WGS-R/DDT-S) are defined as a resistant genotype and susceptible phenotype. VMEs (i.e. false negatives; WGS-S/DDT-R) are defined as a susceptible genotype and resistant phenotype. To detect AMR gene mutations, PointFinder tool for ResFinder and Point mutations tool for AMRFinder results were also included in concordance analyses.

## RESULTS AND DISCUSSION

The main aim of this study was to characterize a collection of Gram-negative CAUTI-associated bacteria (n = 132 isolates) isolated in Egypt to investigate AMR genotype–phenotype concordance in relation to gene data available in widely used, public databases. We also sought to understand the genomic diversity of the dominant species contributing to CAUTIs in an Egyptian clinical setting.

### Genome characterization

Summary statistics for the 132 high-quality genomes assembled in this study can be found in **Supplementary Table 1**. All genomes were assigned to the class *Gammaproteobacteria*, with the exception of Me47, which was found to represent a novel member of the family *Alcaligenaceae* (**Figure (1A)**). Among our isolates, UPEC dominated (50.7 %, 67/132), followed by *Pseudomonas aeruginosa* (10.6 %, 14/132), *K. pneumoniae* (8.3 %, 11/132), other uropathogens (6.8 %, 9/132), *Proteus mirabilis* (6.8 %, n = 9/132), *Providencia* spp. (6 %, n = 8/132), *Enterobacter hormaechei* (3.7 %, n = 5/132), *Alcaligenes* spp. (3 %, n = 4/132), *Serratia* spp. (2.2 %, n = 3/132) and *Pseudomonas vlassakiae* (1.51 %, n = 2/132). UPEC was previously reported as the main cause of UTIs for adults and neonates in Egypt, followed by other *Enterobacterales* or Gram-positive bacteria ^46-49^. *Providencia* – once considered a rare pathogen – is increasingly recognized as an opportunistic pathogen capable of causing serious UTIs ^50^; we found it among the main taxa represented in our study.

**Figure 1:**
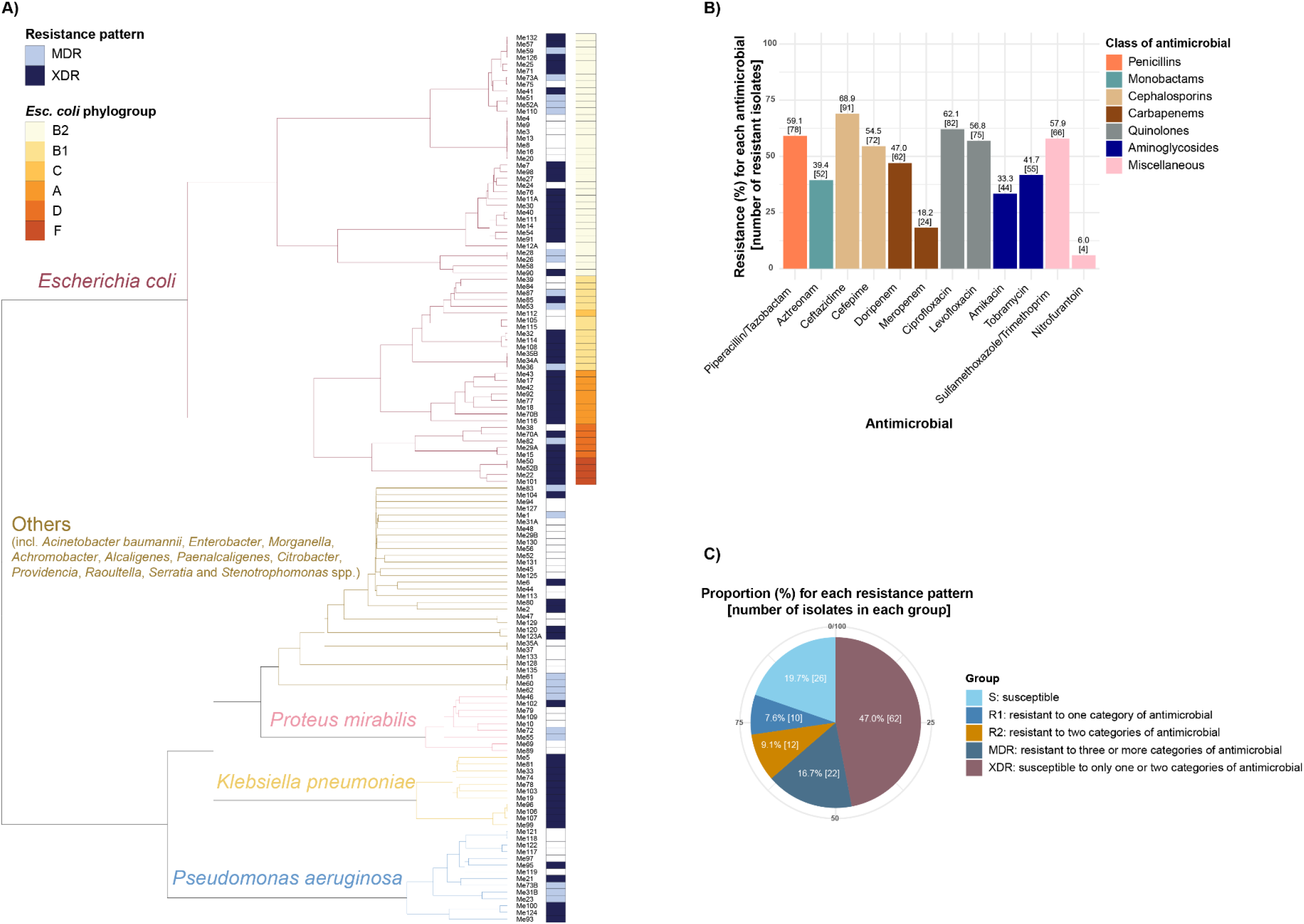
AMR profiles of uropathogens (n = 132 isolates) included in this study. A) Dendrogram generated from comparison of sourmash signatures for uropathogens showing their species affiliations and whether they are MDR or XDR. Phylogroup affiliations are given for *Esc. coli* isolates. B) Resistance summary for the 132 uropathogens using seven different antimicrobial classes according to EUCAST guidelines. (C) Data presented using the category designations suggested by ^54, 55^. Penicillins: piperacillin/tazobactam. Monobactams: aztreonam. Cephalosporins: ceftazidime and cefepime. Carbapenems: doripenem and meropenem. Quinolones: ciprofloxacin and levofloxacin. Aminoglycosides: amikacin and tobramycin. Miscellaneous: sulfamethoxazole/trimethoprim and nitrofurantoin.

### Antimicrobial susceptibility of isolates

According to Egyptian Urological guidelines, symptomatic CAUTIs are treated like complicated UTIs with second- or third-generation cephalosporins, while uncomplicated pyelonephritis is treated with a short course of fluoroquinolones as the first-line therapy ^51^. AMR for quinolones (ciprofloxacin, levofloxacin) and cephalosporins (cefepime, ceftazidime) was >64 % among MDR uropathogens in Egyptian clinical settings ^52^. More than half of UPEC isolates (57.8 %) showed high-level ciprofloxacin resistance and *gyrA* mutations were detected in 76.7 % of isolates in a previous study, leading to recommendations for revision of empirical antibiotic treatment of UTIs ^53^.

Many of our isolates were resistant to ceftazidime (68.9 %, 91/132) and ciprofloxacin (62.1 %, 82/132). Few isolates were resistant to nitrofurantoin (5.9 %, 4/67) and meropenem (18.2 %, 24/132) (**Figure (1B)**; **Supplementary Table 2**). The prevalence of susceptible, R1 and R2 categories ^54, 55^ were 19.6 %, 7.6 % and 9.1 % respectively; 16.6 % (n = 22/132) of isolates were MDR, while 46.9 % of isolates (n=62/132) were XDR (**Figure (1C)**). *Ent. hormaechei* (60 %, 3/5) and *K. pneumoniae* (100 %, n = 11) were associated with the highest prevalence of MDR and XDR, respectively.

### AMR determinants

A wide range of AMR determinants for β-lactams, quinolones, and aminoglycosides have been reported in different uropathogens ^56-59^. The main AMR genes encoded in our genomes were predicted using ResFinder, CARD and AMRFinder (**Supplementary Table 3**). New AMR variants are reported here for uropathogens in Egypt: *OXA-486, OXA-488, OXA-905, IMP-43, PDC-35, PDC-45*, and *PDC-201* for *Pse. aeruginosa*, and *TEM-176* and *TEM-190* for *Esc. coli*.

WGS-based diagnostics may soon take over phenotypic testing for surveillance purposes, particularly in situations where the low error rate has minimal consequences ^60^. Comparison of WGS data and DDT results (with respect to predicted AMR genes and empirical resistance phenotypes) yielded a total concordance of 91.1 %, 85.7 % and 80.3 % for ResFinder, CARD and AMRFinder, respectively. The highest concordance was recorded for *Alcaligenes* (n = 4) species: 97.7 %, AMRFinder; 97.2 %, ResFinder; 100 % CARD (**Supplementary Table 4**).

Variations in discordance were detected for different databases, where the MEs (WGS-R/DDT-S) were 3.9 %, 11.9 % and 15.7 %, while VMEs (WGS-S/DDT-R) were 4.9 %, 2.2 % and 3.8% for ResFinder, CARD, and AMRFinder, respectively (**Figure (2)**). The use of an ineffective therapeutic agent in treatment due to a VME could result in treatment failure, while an ME may restrict therapeutic choices and create complications in the treatment process. The more severe impact of VMEs is evident in FDA regulations for the approval of diagnostic tests, where the FDA mandates that VMEs be kept below 1.5 % and MEs below 3 % for the approval of a new AMR diagnostic test or device ^61^.

**Figure 2:**
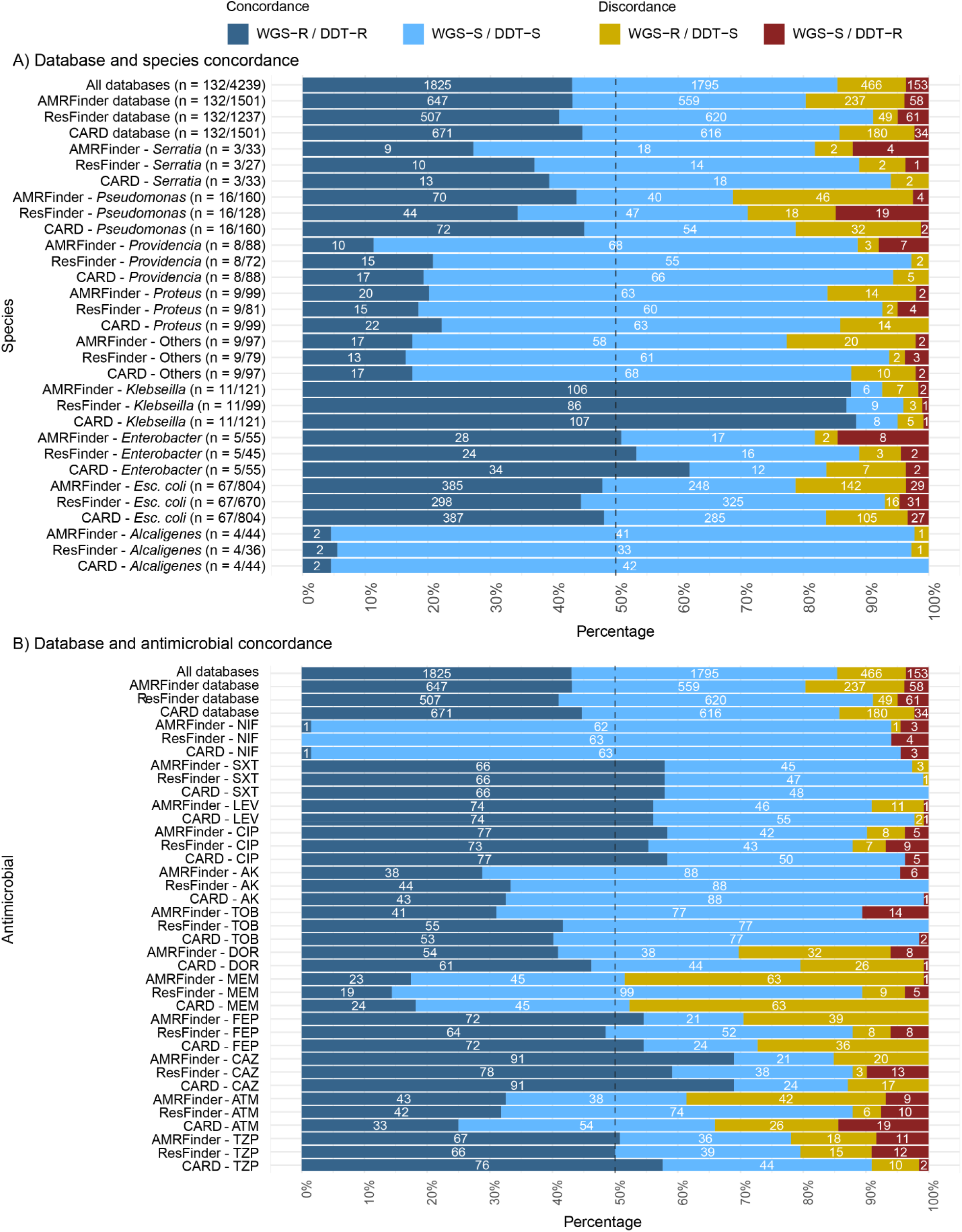
AMR genotype–phenotype discordance summarized for the 132 isolates included in this study. (A, B) Percentage and number of genotype–phenotype concordances and discordances are shown, when comparing WGS data and DDT results using ResFinder, CARD, and AMRFinder. The bars are coloured according to Concordance or Discordance (refer to colour legend at the top of the figure). White numbers on top of the bars correspond to the respective number of results. (A) Species-level data, (B) antimicrobial-level data. Numbers in parentheses in (A) correspond to the number of isolates per species / the number of isolate-antimicrobial combinations. (A) Others: *Paenalcaligenes suwonensis, Klebsiella ornithinolytica, Citrobacter portucalensis, Leclercia adecarboxylata, Morganella morganii, Stenotrophomonas maltophilia, Acinetobacter baumannii*, and *Achromobacter xylosoxidans*. (B) Nitrofurantoin concordance was analyzed only via CARD. Levofloxacin and doripenem were not analyzed with ResFinder/PointFinder. Piperacillin/tazobactam (TZP), aztreonam (ATM), ceftazidime (CAZ), cefepime (FEP), doripenem (DOR), meropenem (MEM), ciprofloxacin (CIP), levofloxacin (LEV), amikacin (AK), tobramycin (TOB), sulfamethoxazole/trimethoprim (SXT), and nitrofurantoin (NIF). Major errors (ME) are WGS-R/DDT-S, while very major error (VME) are WGS-S/DDT-R.

Concordance for aminoglycosides, quinolones, and miscellaneous antibiotics exceeded 89.3 % in all databases. Meropenem was associated with highest discordance in AMRFinder (48.4 %) and CARD (47.7 %), but showed 10.6 % discordance in ResFinder. The range of total discordance of β-lactam antibiotics for the three databases was 10.6–20.4 % ResFinder, 9–47.7 % CARD and 15.1–48.4 % AMRFinder (**Supplementary Table 5**).

In a comparison of CARD and ResFinder for two global collections and a total of 2,587 isolates comprising five pathogens of medical importance, ME rates were higher in CARD (42 %) than ResFinder (25 %). However, CARD showed almost no VMEs (1.1 %) compared with ResFinder (4.4 %) ^37^. In comparison, another report showed an overall concordance of 91 % for ResFinder while ME and VME rates were 6.2 % and 2.1 %, respectively, for 488 different isolates, where *Pse. aeruginosa* had the highest percentage of discordant results (44.4 %) ^44^. In agreement with previous studies ^16, 44, 60^, *Pseudomonas* showed the highest discordance here, and across the three databases used in our study: 31.2 % for AMRFinder, 28.9 % for ResFinder, and 21.2 % for CARD.

In this study, 34.5 % of MEs tested phenotypically “*susceptible, increased exposure*” and were therefore considered as not resistant due to the EUCAST update of susceptibility definitions 2019 “*by the pooling of S and I isolates together into one category*” ^62^. Notably in another study, 22.7 % of MEs tested phenotypically as “*susceptible at higher exposure*” in 234 non-duplicate *Esc. coli* strains ^45^. Discordant bioinformatic predictions of AMR from WGS have previously been shown through analysis of seven clinical isolates with three different databases (ResFinder, CARD and ARG-ANNOT) and where the overall ME “false positives” and VME “false negatives” were 20 % and 14 %, respectively ^63^. In the current study, the overall MEs and VME in three databases were 10.9 % and 3.6 %, respectively. The reduction in MEs and VMEs compared to earlier study ^63^ may in part reflect improvements in curation and coverage of AMR genes through regular database updates over time. The high MEs rate may be also due to low expression levels of enzymes, insufficient understanding of gene regulation, mutations in intergenic regions and *de novo* rRNA mutations.

ResFinder/PointFinder (overall concordance, 91.1 %) showed better *in silico* AMR prediction than CARD (85.7 %) and AMRFinder with Point mutations (80.3 %) (**Figure (2)**). PointFinder tackles chromosome mutation-mediated resistance only in *Esc. coli* and *Klebsiella*. A combined AMR prediction from ResFinder and PointFinder for ciprofloxacin was shown to result in improved prediction performance and a reduced VME rate ^37^.

For accurate *in silico* AMR predictions, our understanding of phenotypic–genotypic correlations must be improved to reach FDA requirements for clinical microbiology diagnostic testing. Excluding genes associated with efflux pump mechanisms from detected AMR genes has been recommended ^37^. Identity thresholds for AMR gene detection should be ≥ 90 % for different databases ^63^. Point mutations leading to AMR should be considered alongside acquired resistance determinants, especially if they are reported from specific clinical sources such as urine, as the most common site for discordance was urine for both *Enterobacterales* and *Pse. aeruginosa*, and mutations in *gyrA* are associated with quinolone resistance of *Enterobacteriaceae* in urine ^64-66^. Databases should replace general terminology (β-lactams, quinolones, etc.) of antimicrobial class as a prediction to confer resistance to all members of the same antimicrobial class, so *in silico* AMR prediction should be tailored to antibiotics themselves rather than antibiotic classes to avoid MEs. For example, the aminoglycoside gene *aac(6*_′_*)-1* leads to amikacin and tobramycin resistance, while *aac(3)-IIa* leads to gentamicin and tobramycin resistance ^67^. This was reported recently as poor correlation (85.9 %) for aminoglycosides between EUCAST and WGS data using ARG-ANNOT and CARD databases, and the authors developed a phenotype-based algorithm to reflect mechanisms responsible for the corresponding aminoglycoside resistance phenotypes ^68^.

Curation of *in silico* databases is required. Some AMR genes listed in databases are not truly responsible for AMR. For example, the *crpP* gene identified by ResFinder and AMRFinder was previously known as a novel ciprofloxacin-modifying enzyme ^69^. However, recent studies showed that *crpP* presence is not always associated with ciprofloxacin resistance ^70-72^. In this study *crpP* was responsible for 7/8 MEs generated by AMRFinder and all seven MEs generated by ResFinder when analysing ciprofloxacin resistance. *crpP* was detected in only one strain (Me73B) resistant to ciprofloxacin and was the only quinolone resistance determinant detected by ResFinder for that strain. However, AMRFinder also detected a point mutation in *gyrA*, T83I, which most likely is the true explanation of ciprofloxacin resistance in Me73B.

Variations between databases can also be found. For example, the AMR phenotype associated with the gene *armA* by ResFinder was amikacin and tobramycin, while AMRFinder predicted resistance to gentamicin. Moreover, the phenotypic prediction for *aac(6’)-Ic* in ResFinder and CARD is resistance to amikacin and tobramycin, while AMRFinder only provides aminoglycoside as subclass.

The *smeF* gene in *Stenotrophomonas maltophilia* is responsible for quinolones resistance ^73^ and was detected by all databases. However, its classification varied: *smeF* was excluded from both CARD and ResFinder as it is considered an efflux pump gene, whereas AMRFinder included it as a resistance determinant for quinolones. This discrepancy contributed to the difference in MEs between ResFinder and AMRFinder for ciprofloxacin in our dataset, with ResFinder showing seven MEs and AMRFinder eight MEs (**Figure (2)**). Although both ResFinder and AMRFinder identified the gene from the exact same contig region with identical coverage, the predicted gene names and associated resistance prediction differed in isolate Me21. ResFinder annotated the gene as *aac(6’)-Ib3*, predicting resistance to amikacin and tobramycin, while AMRFinder identified it as *aac(6’)-Ib*, predicting resistance to gentamicin.

### Detailed analyses of UPEC isolates

Pangenomic approaches highlight genetic diversity, revealing adaptations, virulence factors and resistance traits among collections of genomes. Several pangenome studies of UPEC strains have been performed for both global collections ^74^ and individual countries, including Ireland and Bangladesh ^75, 76^, with the smallest pangenome consisting of at least 16,797 genes in total, the number of core genes ranging from 2,926 to 2,945, and the number of unique genes ranging from 2,900 to 15,266. The UPEC pangenome (n = 67 isolates) here consisted of 10,453 genes: 3,136 (30 %) core; 6,303 (60.3 %) accessory; 1014 (9.7 %) unique.

UPEC isolates from phylogroup B2 were responsible for most of the reported UTIs (n = 36/67, 53.7 %) **(Figure (3)**), in agreement with a previous Egyptian study ^77^. We identified 29 different serotypes (**Supplementary Table 1**), with phyologroup B2 serotypes O25:H4 (n = 12/67, 17.9 %) and O75:H5 (n = 11/67, 16.4 %) most common. Although phylogroup B2 strains are uncommon among the commensal microbiota, they are highly virulent when present; B2 strains have been shown to persist in the intestinal microbiota of infants ^78^, while also being associated with wildlife and livestock ^79^, and herbivorous and omnivorous mammals ^80^. A previous Egyptian study showed the most prevalent phylogroup in the country was A followed by B2 ^81^. Other phylogroups detected by us were B1 (19.4 %), A (11.9 %), D (7.4 %), F (5.9 %) and C (1.4 %).

The principal virulence genes associated with all UPEC strains (n = 67) were type 1 fimbriae (*fimH*), outer membrane protein (*ompA*) and enterobactin (*entS*). Curli fibers (*csgA* 98 %, n = 66), common pilus (*ecp* 95 %, n = 64), nutritional and metabolic factors (*fyuA* 79 %, n = 53, *chuA* 67 %, n = 45), K1 capsule type (*kpsM* 66 %, n = 44), aerobactin (*iutA* 60 %, n = 40), P fimbriae (*papC* 52 %, n = 35, *papG* 33 %, n = 22), α-hemolysin and iron siderophores (*hlyA* and *iroN* 24 %, n = 16), exotoxins (*cnf1* 13 %, n = 9), and S fimbriae and F1C fimbriae (*sfa* and *foc* 6 %, n = 4) were also represented (**Figure (3)**; **Supplementary Table 6**). *fimH* and *sfaX* were present in all phylogroups; all other *sfa* genes were only associated with phylogroup B2. *entD* was only in phylogroup B1. *cnf-1* was present in all phylogroup B2 isolates and only one isolate (Me87) from B1.

**Figure 3:**
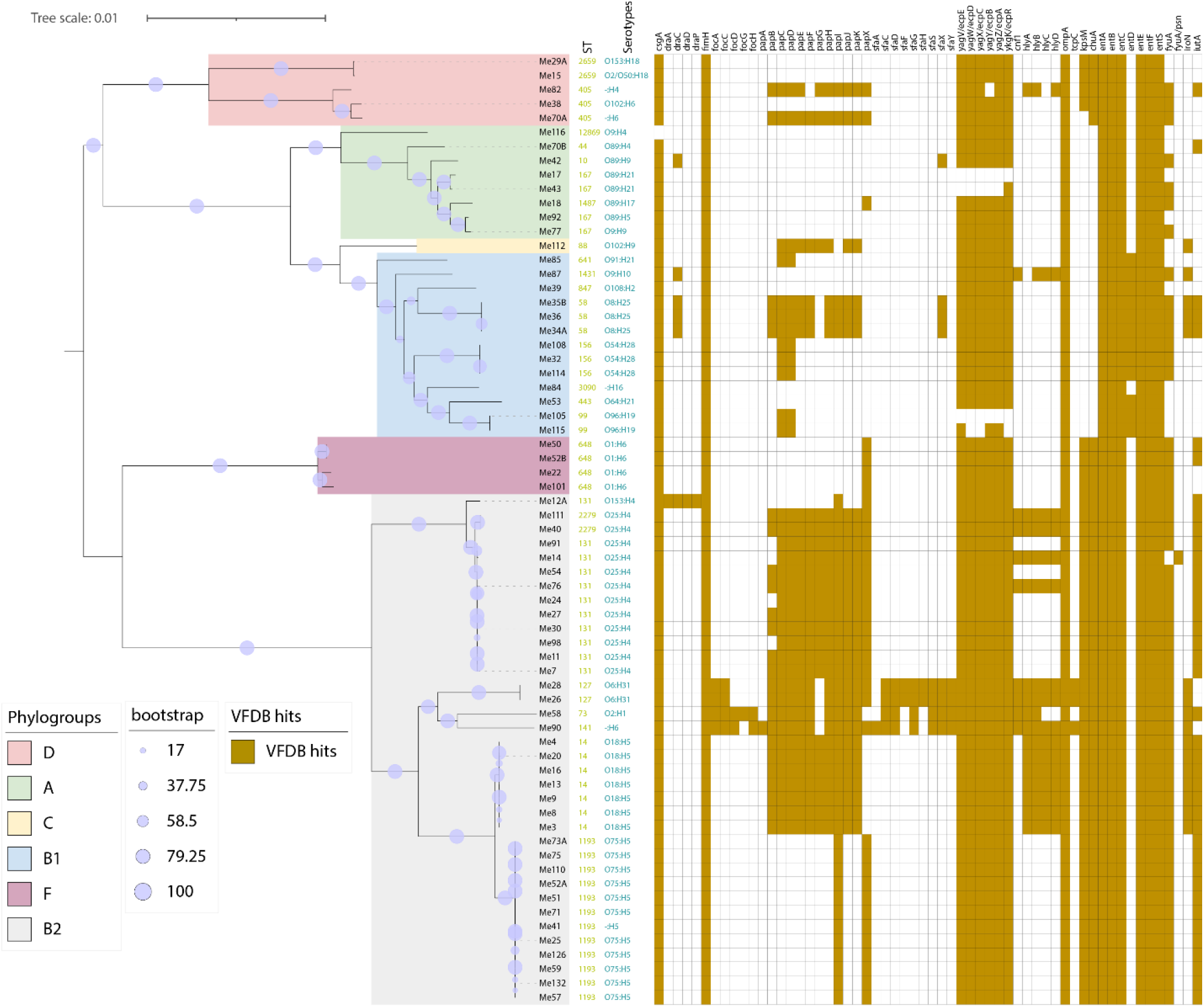
Phylogenetic tree for 67 *Esc. coli* isolates constructed using pangenome analysis of core genes, via panaroo and RaxML. UPEC strains are grouped into clusters (A, B1, B2, C, D, F) depending on EzClermont phylogroups. The maximum-likelihood tree was annotated using iTOL, with bootstrap values expressed as a percentage of 1000 replicates. Scale bar, nucleotide substitutions per site. Virulence genes were identified through BLASTp with VFDB based on coverage >= 90 % and identity >=70 %.

*fim, ecp* and *csg* gene families have previously been found to be core to the UPEC pangenome ^76^. In contrast, a PCR-based study from Mansoura reported a much higher prevalence of afimbrial adhesins such as *afa/dra* (14 %) and S-fimbriae (*sfaS*, 60.6 %) ^82^, which were largely absent from our dataset. Additionally, our data show a higher prevalence of P fimbriae genes (*papG*, 33 %; *papC*, 52 %) compared to ^82^, where genes were found only in only 21.3 % and 49.3 % of isolates, respectively. Similarly, our detection of iron acquisition genes *iutA* (60 %) and *chuA* (67 %) exceeds their reported values of 34.6 % and 54 % ^82^. Notably, *cnf1* (cytotoxic necrotizing factor 1) was much rarer in our dataset (13 %) compared to 42 % in ^82^, suggesting potential differences in strain pathogenicity. Overall, our results highlight a strong presence of core adhesion and iron acquisition systems in UPEC strains, with notable variations in toxin-associated genes compared to previous studies. These findings emphasize the genetic diversity of UPEC virulence factors and the importance of comparing detection methods when assessing pathogenic potential.

The β-lactam resistance genes *bla*_OXA-1_ and *bla*_TEM-1_ were present in all phylogroups, *bla*_TEM-176_ was restricted to phylogroup B1, while other *bla*_TEM_ variants were present in only phylogroup A. *dha* gene variants were restricted to phylogroup B2, while *bla*_NDM-5_ was absent from B2. *bla*_CTX-M-15_ was present in all phylogroups, while *bla*_CTX-M-14_ and *bla*_CTX-M-16_ were present in phylogroup F only. In Egypt, *bla*_CTX-M-15_ is the most predominant ESBL-producing type among uropathogenic and diarrheagenic *Esc. coli* strains ^83^. In terms of quinolone resistance, *qnrB4* was present in phylogroup B2 only, while *qnrS1* was present in all phylogroups. Mutations of *gyrA* and *parC* genes were present in all phylogroups. Genes of aminoglycoside-modifying enzymes were present in all phylogroups.

UPEC isolates were most susceptible to nitrofurantoin (3 % resistance). Nitrofurantoin was reported as a drug of choice for treating UTIs in previous Egyptian studies ^84, 85^, but it has therapeutic limitations due to side-effects in certain patient groups ^86^. *In silico* AMR prediction using CARD showed 95.5 % concordance with phenotypic results (**Figure (2)**). In a recent Egyptian study, the resistance level to nitrofurantoin was 51 % in *Klebsiella* spp., but it remains effective for *Esc. coli* UTIs in males and females (92 % sensitivity). In comparison, *Esc. coli* and *Klebsiella* spp. showed elevated resistance rates to cephalosporins: 75 % and 81 %, respectively ^87^.

## Conclusion

Our study provides crucial insights into the genomic and AMR landscape of CAUTI-associated uropathogens in Egypt, highlighting the importance of understanding AMR phenotype–genotype discordance in this setting. These findings reinforce the need for integrated phenotypic and genomic surveillance strategies to improve AMR diagnostics, guide treatment decisions, and inform infection control policies in resource-limited settings.

## Abbreviations

AMR: antimicrobial resistance
CARD: Comprehensive Antibiotic Resistance Database
CAUTI: catheter-associated urinary tract infection
DDT: disc diffusion test
ESBL: extended spectrum β-lactamase
HAI: healthcare-associated infection
ICU: intensive care unit
MDR: multidrug-resistant
ME: major error
MLST: multilocus sequence type
VME: very major error
UPEC: uropathogenic *Escherichia coli*
UTI: urinary tract infection
XDR: extensively drug-resistant

## Data availability

The sequence data included in the study are available under BioProject PRJNA1208035.

## ACKNOWLEDGEMENTS

We would like to Dr Essam Elsawy and staff of Urology and Nephrology Centre, Mansoura, Egypt for providing the clinical isolates used in this study.

ME – did all phenotypic work; extracted DNA for sequencing; all bioinformatics associated with clinical isolates; characterized the AMR and virulence genes encoded by the isolates and their plasmids; MLST analysis and summary; interpreted virulence and AMR data; pangenome analysis. JCT – pangenome analysis and associated supervision. NH, DN – library preparation and genome sequencing of some clinical isolates; LH – sourmash analyses; supervised the study. ME, JCT and LH wrote the original version of the manuscript, and all authors approved the final version.

## TRANSPARENCY DECLARATIONS

The authors declare that there are no conflicts of interest.

